# St2cell: Reconstruction of in situ single-cell spatial transcriptomics by integrating high-resolution histological image

**DOI:** 10.1101/2022.10.13.512059

**Authors:** Siyu Hou, Kuan Tian, Sen Yang, Jinxi Xiang, Wei Yang, Jun Zhang, Xiao Han

## Abstract

Spatially resolved transcriptomics (SRT) has greatly expanded our understanding of the spatial patterns of gene expression in histological tissue sections. However, most currently available platforms could not provide in situ single-cell spatial transcriptomics, limiting their biological applications. Here, to in silico reconstruct SRT at the single-cell resolution, we propose St2cell which combines deep learning-based frameworks with a novel convex quadratic programming (CQP)-based model. St2cell can thoroughly leverage information in high-resolution (HR) histological images, enabling the accurate segmentation of in situ single cells and identification of their transcriptomics. Applying St2cell on various SRT datasets, we demonstrated the reliability of reconstructed transcriptomics. The single-cell resolution provided by our proposed method greatly promoted the detection of elaborate spatial architectures and further facilitated the integration with single-cell RNA-sequencing data. Moreover, in a breast cancer tissue, St2cell identified general spatial structures and co-occurrence patterns of cell types in the tumor microenvironment. St2cell is also computationally efficient and easily accessible, making it a promising tool for SRT studies.

## Main

Rapidly emerging SRT technologies are enabling the whole-transcriptome measurement of spatial locations on an unprecedented scale, which provides great opportunities to dissect the spatial architecture and functional organization of multicellular tissues from various biological contexts^1–3^. However, the widely available platforms which adopt in situ RNA capturing (ISC) technologies (such as ST^4^ and 10x Visium) measure the gene expression profile at the basic spatial units of spot^1, 5–7^, which may hardly overlap with individual cells and usually have relatively lower spatial resolutions compared with the scale of one single cell (for example, typically 3-30 cells contained in one spot from 10x Visium^4, 8^). This weakness of the “non-single-cell” characteristic seriously limits their potential values in biological applications. Therefore, it is an essential challenge to enhance the spatial resolution, and especially obtain the cellular level spatial transcriptomics.

Though a few relevant SRT resolution enhancement methods have been developed recently, these methods tend to perform only limited enhancement. As an example, BayesSpace^9^ uses statistical model combined with information from spatial neighborhoods to enhance the resolution of SRT. However, this enhancement method based on the SRT data itself can only increase the resolution to 6-9 times that of the original data. On the other hand, another class of approaches focuses on the association between gene expression and anatomical features (or histological images). For instance, Xfuse^10^ has produced super-resolved spatial transcriptomes by introducing HR image information paired with spatial transcriptome data. However, these types of approaches sacrifice the time efficiency due to pixel-by-pixel prediction of gene expression. More importantly, both types of methods ignore the location of cells in the tissue and merely obtain pseudo-cellular level resolution. To address this algorithmic hurdle in enhancing the spatial resolution of SRT as well as overcome the limitations of most currently available platforms, we develop a new framework to take advantages of both information from SRT data and co-registered HR histological image.

Our method, called St2cell, a computational framework that fuses HR histological images and SRT data to infer the in situ single-cell transcriptomics. Indeed, HR histological images contain a wealth of information. Many recent studies have revealed extensive correlations between image features and gene expression levels as well as histopathology^11, 12^, and there are even studies that can predict pathology-related gene expression levels in a weakly supervised manner using only HR pathology images^13^. All these suggest that combining SRT and HR histology images can help to reveal underappreciated information. Here, we innovatively use deep learning models to localize cells and extract cell features, and perform in situ single-cell transcriptomics inference using a CQP-based model. With the introduction of deep models, St2cell can effectively utilize SRT information from spatial neighborhoods and accurately distinguish between cellular and tissue contexts. We extensively applied it to several publicly available SRT datasets and conducted different application analyses. 1) We first achieved in situ single-cell gene expression profile reconstruction using St2cell on Visium FFPE adult mouse brain sections, demonstrating that our approach is capable of reconstructing intra- and inter-spot expression levels at single-cell resolution and genome-wide. 2) We compared our approach with other methods through a spatial domain detection task and two simulated experiments on ST HER2-positive tumors datasets with manual annotation and confirmed the accuracy of the St2cell reconstruction results. 3) We analyzed a set of Visium mouse brain with matched single-nucleus RNA sequencing (snRNA-seq) data and found that St2cell can help map cell distribution. 4) Case studies on Visium breast cancer dataset further identify the advantages of St2cell in facilitating the detection of cell types and recovery of the distribution of tumor micro-environmental markers.

Collectively, we show that St2cell can infer in situ single-cell transcriptomics, enhance spatial resolution and contribute to the downstream analysis, allowing us to characterize the molecular organization of tissues and organs more finely. These results demonstrate that our method is a promising tool that can effectively exploit the rich information contained in the HR histological image data and integrate it with the spatial gene expression profiling.

## Results

### Overview of St2cell workflow

Our St2cell framework consists of three main components, including cell localization and cell feature extraction from HR histological images, and further inference of the in situ single-cell transcriptomics through the incorporated analysis with the measured spatial transcriptomics (Methods; Fig. 1). In situ single-cell localization is achieved by a heatmap regression-based convolutional neural network (CNN) framework, which can be used in conjunction with the known radius and coordinates of the spot in SRT for determining which spot each cell belongs to. Next, we deploy a contrastive learning-based image feature extraction model that assigns soft labels representing cell categories (treated as meta-cell types) to each cell by automatically learning the morphological differences between cells. Combined with the spatial locations obtained in the previous step, we can further obtain the meta-cellular composition of each spatial spot. Finally, we combine the measured SRT profiles with the above cell features and cell locations together to model the inference of the gene expression levels of cells as a quadratic programming problem with constraints, and design a CQP-based model to decouple the mixed single-cell transcriptomics in spots by minimizing the error between the reconstructed and measured gene expressions (Methods). We argue that the way St2cell integrating HR images is essential for effective recovery of the in situ single-cell transcriptomics, and have experimentally validated that St2cell can accurately reconstruct SRT at the single-cell resolution in silico.

**Figure 1.**
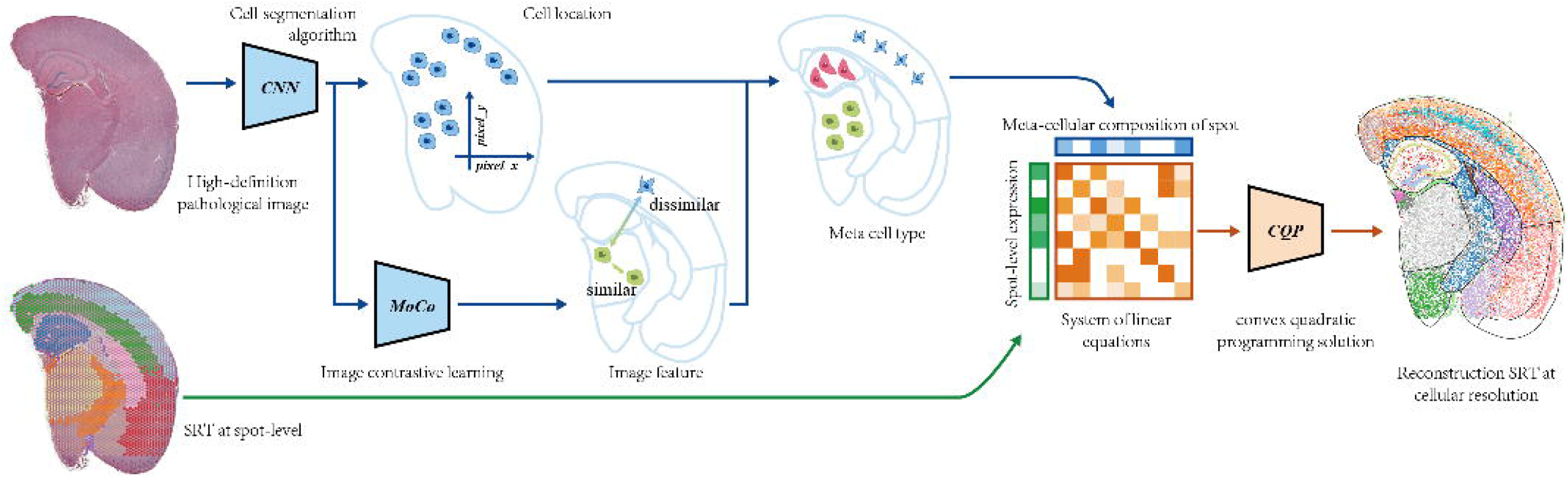
Overview of the St2cell. The input to St2cell is low-resolved SRT and co-registered HR histological images. St2cell draws on deep learning models to query cell locations and extract cell features and employs a CQP-based model to estimate gene expression levels for each cell. St2cell fuses data from these two distinct modalities to enable in situ single-cell transcriptomics reconstruction that facilitates downstream analysis.

### St2Cell accurately identifies single-cell spatial patterning of mouse brain

To demonstrate the ability of St2cell to recover the in situ single-cell expression profiles, we first applied the model to data from the adult mouse brain, of which the well-defined spatial architecture is function-related and could serve as a canonical surrogate for evaluating the precision of reconstructed transcriptomics^14–16^ (Fig. 2A). We manually annotated the tissue section assayed into regions as the cortex (CTX), hippocampus (HPC), white matter (WM), thalamus (THA), hypothalamus (HYP), amygdala (AMY), prefrontal cortex (PFC) and striatum (STR), as suggested by previous studies^17, 18^. St2cell was deployed to demarcate the location of all single cells from the matched HR histological image, and then reconstruct their gene expression profiles through integrative analysis with the measured transcriptomics of spatial spots. The results demonstrated that St2cell can successfully boost the spatial resolution to the cellular level, as well as reliably retain the histological architectures.

**Figure 2.**
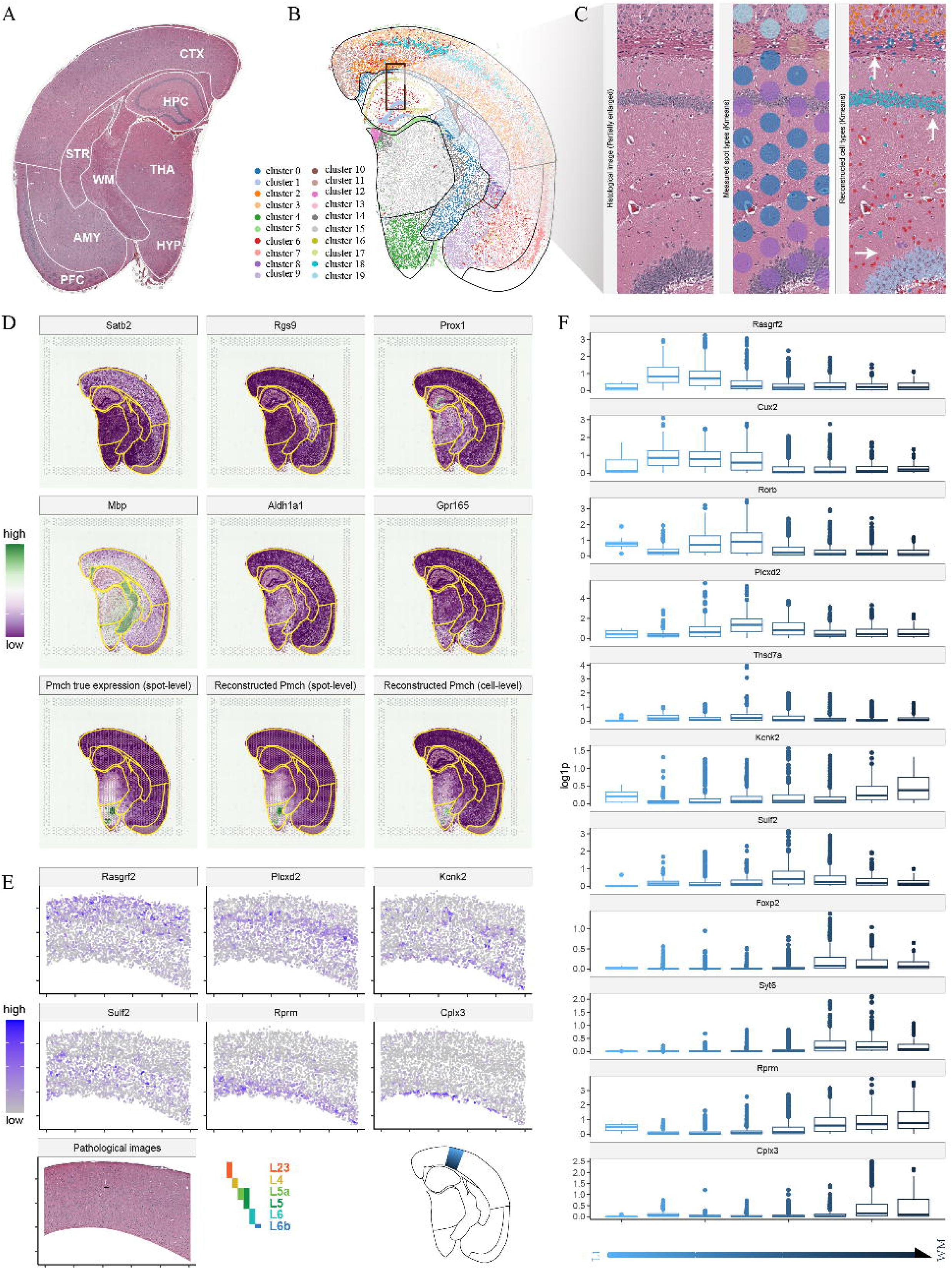
St2cell identifies single-cell spatial patterns of mouse brain. (A) Co-registered HR histological images of adult mouse brain and its manually annotated tissue regions. (B) Clusters of St2cell-reconstructed in situ single-cell gene expression profiles. (C) The partial enlarged figure of areas including the CTX, WM, HPC. Left panel: original FFPE H&E stained histological image. Middle panel: The size and position of the spots, with different colors representing different categories after clustering of the measured spots’ transcriptomics. Right panel: The locations and types of cells inferred by St2cell, with different colors representing different categories after clustering of the cellular-level gene expression profiles obtained by our proposed method. (D) Spatial gene expression patterns of regional highly expressed markers. (E) Spatial gene expression patterns of six markers of layer-specific cortex pyramidal cells. (F) Spatial distributions of layer-specific expressed genes at different locations.

Indeed, we divided the in situ single cells into twenty categories that far exceed the number of manually annotated regions in order to test the ability of St2cell to automatically delineate tissue regions and even cell types (Methods). It shows that St2cell is able to delineate tissue regions consistent with manual annotation and can further identify sub-regions with cellular layer-enrichment specificities, such as CTX and HPC (Fig. 2B). To further illustrate the superiority of St2cell, we intercepted a partially enlarged HR image across three regions of CTX, WM and HPC as a demonstration (Fig. 2C left). As shown in Fig. 2B and 2C, the reconstructed single-cell transcriptomics of St2cell provided a higher-resolution cellular map of the tissue with a more distinctive and accurate regional delineation, enabling it to successfully remove the inherent bottlenecks of the original 10x Visium data. For example, the sparse location distribution of captured spots (10x Visium dataset with a spot diameter of 55 μm and a spacing of 100 μm between spots, Fig. 2C middle) could not cover the entire tissue region, and furthermore, each spot contains multiple cells, which mixes together the expression profiles of different cell types. Promisingly, St2cell succeeded in reconstructing the intra- and inter-spot cellular gene expression, obtaining in situ single-cell transcriptomics (Fig. 2C right). The results of St2cell closely matched the brain architecture and divided the cerebral cortex into different subregions meticulously. We observed clear demarcations at the boundaries of CTX, WM and HPC subregions consistent with real histological structures. Furthermore, the cell clustering results of St2cell explicitly distinguished between somatic and dendritic layers (cyan and light blue points) in the HPC region, which could not be discriminated using the measured transcriptomics of spots (purple spot) in contrast.

To consolidate the reliability and interpretability of St2cell results, we next detected regional markers that are highly expressed in the particular annotated domain but are expressed at low levels in the neighboring regions. As an example, we identified Satb2, Rgs9, Prox1, Mbp, Aldh1a1 and Gpr165 in CTX, STR, HPC, WM, THA and HYP regions, respectively, based on St2cell results, which are also confirmed by previous studies^19–23^. The results (Fig. 2D, Supplementary Fig. S1) show that the gene expression levels reconstructed by St2cell are highly expressed in these specific anatomical regions, which agree well with known tissue structures. In addition, we used the example of Pmch, another detected spatially variable gene (SVG) that is significantly highly expressed in the HYP region, and showed that the St2cell results well fit the spatial pattern of the measured spot-level gene expression profile and can further provide the accurate location of cells with high Pmch expression.

Finally, we sought to explore whether St2Cell could achieve the fine-grained spatial mapping of gene expression. We zoomed in and elaborately cropped the region of the cortex containing L23, L4, L5a, L5, L6 and L6b, inspecting the spatial gene expression patterns of six layer-specific cortex pyramidal cells markers – Rasgrf2 (L23), Plcxd2 (L4), Kcnk2 (L5a), Sulf2 (L5), Rprm (L6), and Cplx3 (L6b)^24^. Consequently, the expression distribution of all these markers exhibits a high degree of layer-enrichment specificity, which are also largely congruent with those obtained from in situ hybridization (ISH) data^25^ (Supplementary Fig. S2), allowing alignment with known locations of sub-layers of the cortex. We note that for the extremely narrow L6b region, in particular, St2cell enriched the expression of regionally specific Cplx3 in this pinpoint position, perfectly segmenting the adjacent WM and CTX region (Fig. 2E; Supplementary Fig. S3). Additionally, we extended to unveil the dynamical change of gene expression profiles as the cell location evolved from the shallow layer to the deep of the cortex (Fig. 2F). The well-characterized marker Cux2 was highly expressed in the shallow layer, and reached the peak expression at L23. Rorb and Thsd7a were enriched in the middle L5a layer, while Foxp2, Syt6, Rprm were specifically expressed in deeper layers, such as L6 and L6b.

Taken together, St2cell could convincingly reconstruct the in situ single-cell transcriptomics and the cellular-level spatial atlas with reliable soundness guarantees and would bring about a better understanding of the molecular organization of tissues.

### Benchmark evaluations of St2cell

Next, a spatial domain detection task and two simulation experiments were set up to quantitatively benchmark the performance of St2cell and compare it with other related algorithms, in terms of both reliably identifying the spatial architecture and precisely reconstructing the transcriptomics. We adopted a dataset of HER2-positive tumors (termed “ST BrCa data”) which contains various cell types, and the spatial domains of each sample has been well-annotated by expert pathologists^26^.

The spatial regions in these samples were recognized into seven different categories - “adipose tissue”, “breast glands”, “cancer in situ”, “connective tissue”, “immune infiltrate”, “invasive cancer” and “undetermined” as provided by the original study. We used St2cell as well as four counterparts - Kmeans, Louvain^27^, SpaGCN^3^ and BayesSpace^9^ to detect the spatial domains from the SRT data, evaluating their accuracy with the annotated ones respectively (Methods). Two representative samples – “G2” and “H1” with much more complicated tissue structures than others were selected to demonstrate the method performance. Qualitative visualization shows that the spatial domains detected by St2cell are largely congruent with the annotated regions, while the other algorithms show inferior consistency (Fig. 3A). For example, St2cell precisely identified the region of “immune infiltrate” in both samples. By contrast, in the “H1” section, Louvain labeled “immune infiltrate” and part of “cancer in situ” as one category, while SpaGCN failed to identify the exact boundaries of “immune infiltrate”, resulting in the erroneous inclusion of some “adipose tissue” regions. Furthermore, all methods except St2cell were unable to accurately discern the “immune infiltrate” regions in the G2 samples. Accordingly, quantitative metrics also show that St2cell could significantly outperform all other algorithms in this task, achieving an ARI of 0.278 for section G2 and 0.401 for section H1. In comparison, the ARI of other methods are much lower (0.171 and 0.264 for Kmeans, 0.161 and 0.218 for Louvain, 0.126 and 0.319 for SpaGCN, 0.173 and 0.374 for BayesSpace). Although SpaGCN and BayesSpace utilized histological or adjacency spot information, their performance is not much better than Kmeans and Louvain and are substantially worse than St2cell. Notably, compared to other methods, St2cell not only detects at the spot level but can further delineate the spatial domains at the cellular level, giving boundaries similar to those of manual annotation and providing additional spatial details (Fig. 3A cell-level).

**Figure 3.**
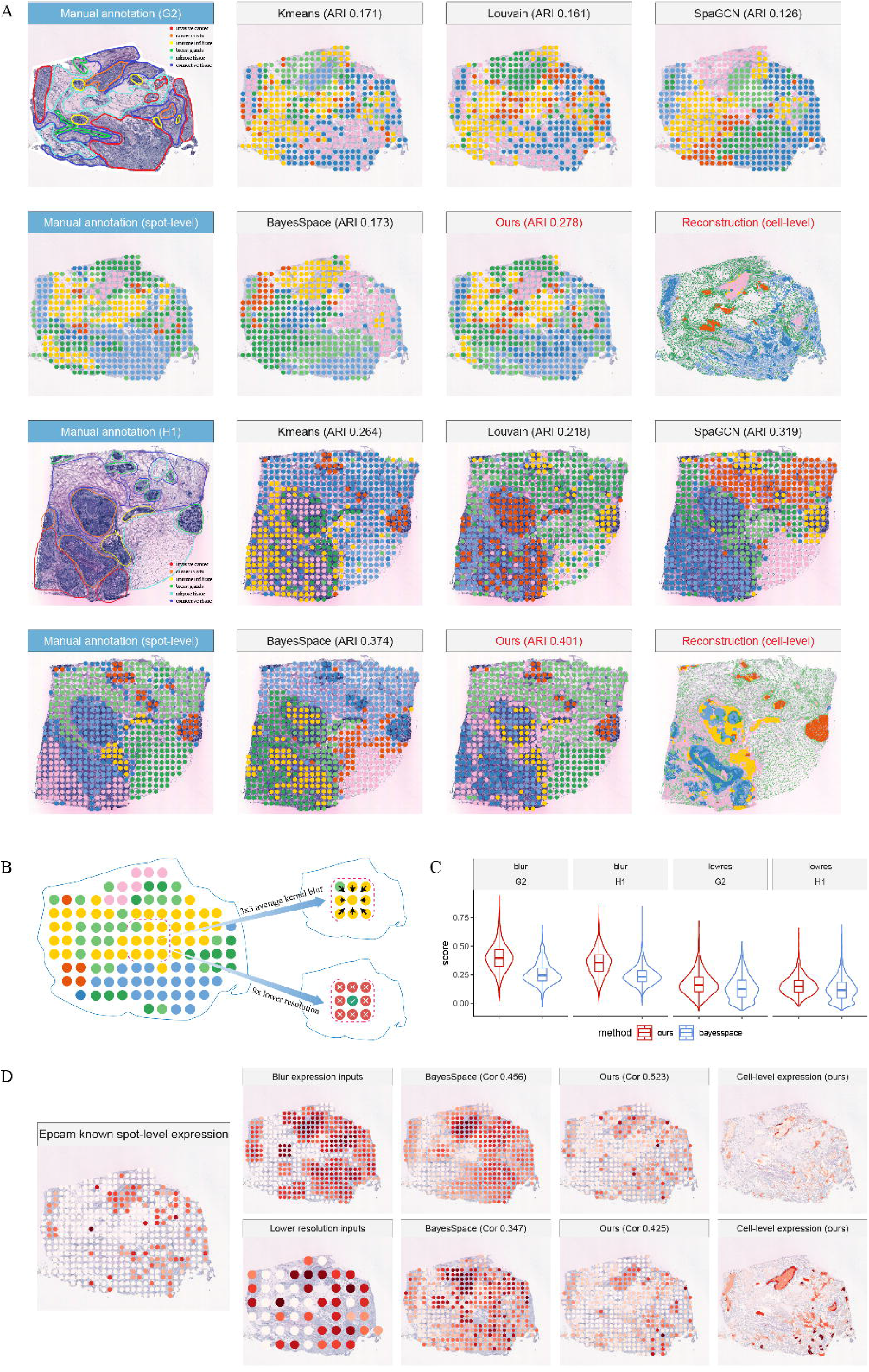
St2cell’s benchmarking and simulation results. (A) Spatial domains detected in the ST breast cancer data. Left: Histological images of sections from the original study with manually labeled regions at different levels. Grids with red captions represent the spatial domains obtained by St2cell. Different colors represent different clusters. (B) Schematic diagram of the two different simulation experiments. (C) Pearson’s correlation values of the 2,000 HVGs expressions reconstructed by St2cell and BayesSpace. (D) Reconstructed Epcam expression level of “G2” by St2cell and BayesSpace under two different simulations.

We then evaluated the capability of St2cell to improve the spatial resolution with high fidelity through two simulation experiments. The blurred spatial gene expression profiles and lower-resolution of captured locations were input to the model respectively, assessing the conformance between the algorithm output and the original data to benchmark the reconstruction precision against BayesSpace (Fig. 3B). In simulation 1, we simulated the contamination of spot’s gene expressions by surrounding spots using ST BrCa data. For each spot of the sample, we averaged the gene expression levels of its surrounding spots using a 3×3 kernel to obtain the blurry expression profiles of the corresponding captured locations. In simulation 2, we dropped 89% of the data, and for every 3×3 captured locations, we keep only the middle one as the spot for the generated data. Using the Pearson correlation coefficients between the reconstructed gene expression profiles of each spot and the original data to compare the performance of St2cell and BayesSpace, we found that St2cell outperformed BayesSpace in terms of yielding substantially higher concordance for the 2,000 highly variable genes (HVGs) as shown in Fig. 3C. Specially, in simulation 1, the median correlations of St2cell were 0.404 and 0.358 and BayesSpace were 0.261 and 0.244 for G2 and H1 samples, respectively; in simulation 2, the median correlations of St2cell were 0.172 and 0.157 and BayesSpace were 0.137 and 0.123. Finally, we selected a cell surface molecule Epcam, known to be highly expressed in breast and epithelial carcinomas^28, 29^, as a concrete example to confirm the reconstruction results of our method. As shown in Fig. 3D, the reconstructed Epcam expressions (also see the results of “H1” in Supplementary Fig. S4) were better coordinated with the ground truth, with the correlations achieved bySt2cell are significantly higher than those by BayesSpace.

Overall, these results suggest that St2cell is a robust method to precisely improve spatial resolution and moreover to comprehensively characterize underlying biological structures from the spatial transcriptomics data.

### St2cell promotes the smooth integration with single-cell (nucleus) transcriptome data

Integrative analysis with scRNA- or snRNA-seq data from the same tissue can potentially facilitate the interpretation of SRT data in terms of matching cell type information to the spatial location^30–32^. However, current methods fail to directly mitigate the issue of mixed cell types induced by the limited resolution of SRT. In contrast, the in situ single-cell transcriptomics manufactured by St2cell offers a more explicit way to transfer information from scRNA- or snRNA-seq data. Here, to validate this versatility of the St2cell, we analyzed another mouse brain dataset containing multiple regions generated from the telencephalon and diencephalonl^18^. Unlike our previous FFPE adult mouse brain data, this dataset further contains matched single-nucleus data along with the spatial RNA-seq profiles from five adjacent mouse brain sections. Each section includes about 3,000 spots and 31,000 genes.

We first noted that compared with our other experimental datasets, the paired histological images of this data were relatively low-quality, making it more challenging to distinguish cell morphology for St2cell. As in the previous operation, we clustered the reconstructed in situ single-cell transcriptomics into twenty categories and labeled the clusters with different colors (Fig. 4A). Remarkably, the results show that St2cell can still robustly perform reconstruction and the clustered categories as a whole obey the results of the previous regional delineation of brain tissue (see also Supplementary Fig. S5).

**Figure 4.**
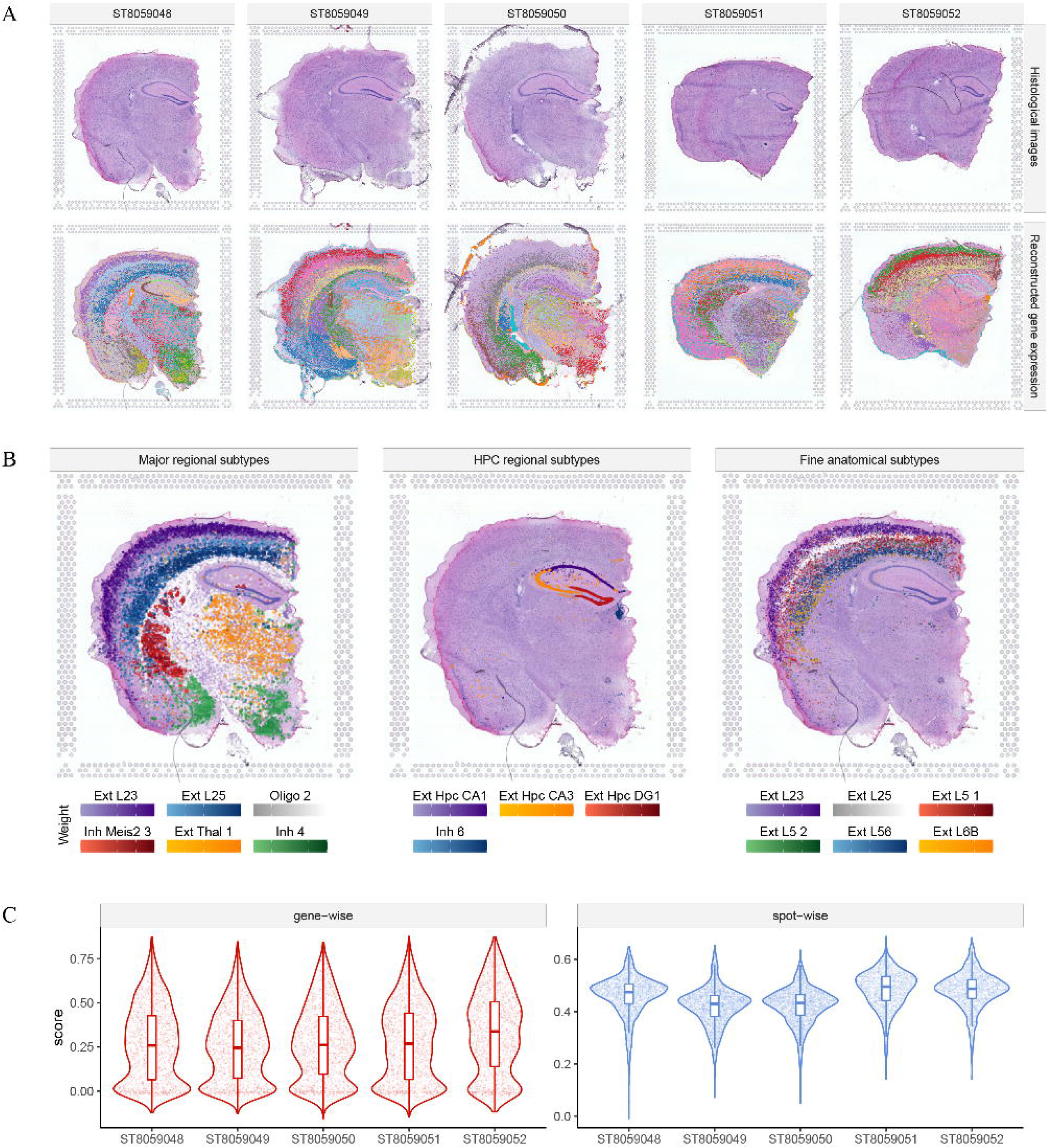
Application of St2cell to adjacent mouse brain sections with matched single-nucleus and Visium spatial RNA-seq profiles. (A) Clustering results of cellular resolved spatial gene expression profiles reconstructed by St2cell. (B) Distribution and abundance of cell types estimated from the results of St2cell. Left: spatial patterns of major regional subtypes. Middle: spatial patterns of sub-regional cell types. Right: spatial patterns of fine anatomical subtypes. (C) Correlation of the integrated spot profiles with the known measured profiles.

Next, we adopted a domain adaptation algorithm and k-nearest-neighbor regression^30, 33^ to align cells from the snRNA-seq data with spatial single cells output by St2cell, explicitly mapping the spatial distribution of every cell type (Methods). As shown in Fig. 4B, St2cell captured different spatial patterns of cell types, mapping a wide range of regional specific cell types such as excitatory neurons of THA and oligodendrocytes of WM, sub-regional types such as excitatory neurons of CA1, CA3 and DG1 cells of HPC, and sub-regional cell types across diverse cortical layers. It demonstrated that St2cell have the advantages in disentangling measured transcriptomics of spots to cellular resolution and enabling direct and convenient integration of single-cell data on the ISC technology platform as on the ISH technology platform.

Finally, we calculated Spearman correlation coefficients at gene-wise and spot-wise of integrated results. These coefficients can directly reflect the relevance between reconstructed and measured spatial gene expression profiles. As shown in Fig. 3C, the vast majority of the gene expressions obtained from the reconstructions were positively correlated with the measured levels at gene-wise. In particular, for all five samples, the average correlations at spot-wise were over 0.4, indicating that St2cell can effectively portray the differences between spots, laterally indicating that St2cell can help in cell type estimation and spatial domain detection. All these results demonstrated that St2cell reconstructed gene expression at the single-cell level, which indeed facilitated to integrate scRNA or snRNA-seq data, as well as to gain a more comprehensive understanding of the structure of cell type distribution.

In summary, we show that St2cell can considerably break down the barriers between the single-cell-level scRNA- or snRNA-seq-based reference data and the spot-level SRT data, decoupling the integration process of annotating the spatial distribution of cell types.

### St2cell identified the co-occurrence patterns of cell types within the tumor microenvironment

The cellular level spatial atlas provided by St2cell could enlighten researchers on both the spatial distribution patterns of individual cell types and their co-occurrence. To further examine the ability of St2cell to effectively delineate complicated co-location patterns within the tumor microenvironment (TME), we further investigated another breast cancer sample with a richer spatial structure. St2cell was applied to recover the in situ single-cell transcriptomics, and a scRNA-seq dataset from breast cancer with annotated cell type information was then leveraged to infer the cell identity of in situ single cells (Fig. 5A; Methods).

**Figure 5.**
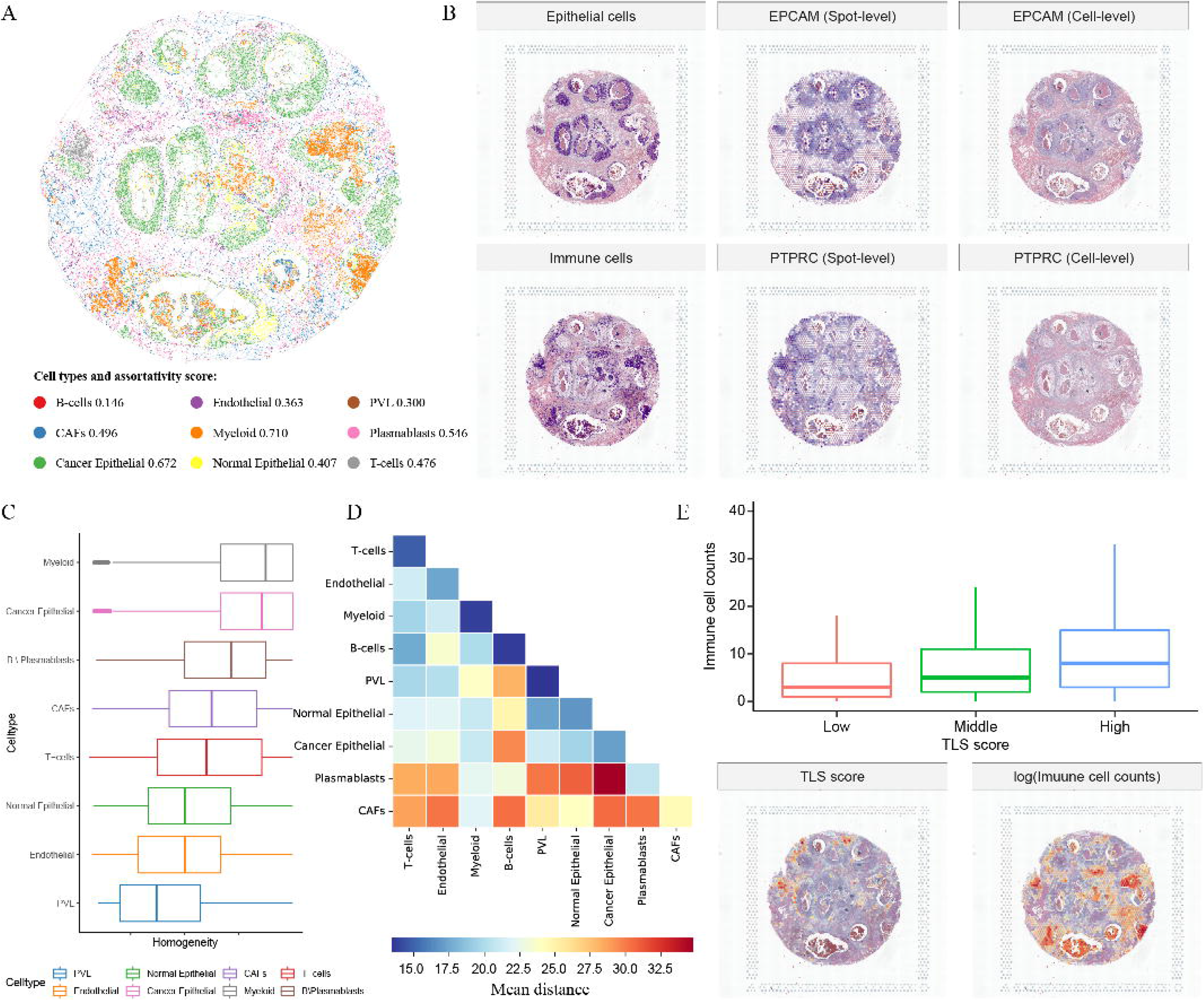
St2cell identified the co-occurrence patterns of cell types within the tumor micro-environment. (A) The spatial graph constructed by Delaunay triangulation based on results of St2cell. Different colors represent different annotation results. (B) Spatial distribution of epithelial and immune cells and the spatial expression patterns of their markers under different levels. The proportion of cells surrounded by the same type of cells, with lower scores indicating more random mixing. (D) Heat map of the mean distance between two contacting cells of different cell types. (E) The relationship between TLS score and immune cell counts.

Checking the expression of cell type-specific markers, we found a great congruence between the inferred cell identities and their corresponding markers (Fig. 5B). For example, cells annotated as epithelial cells were mostly located in spots with high expression of Epcam, and cellular level gene expression profiles reconstructed from St2cell accurately recapitulate this feature. We also visualized the spatial expression pattern of Ptprc, which was in accordance with the locations of immune cells with high specificity. Correspondingly, the districts in which cancer epithelial cells located were characterized by unique histological image features, and both the cancer epithelial cells and immune cells coincided with the tissue region classification by expert physicians (Supplementary Fig. S6). These results indicate that St2cell combined with the integration of scRNA-seq annotations could plausibly recover the cell type identities of in situ single cells.

Incorporating both the spatial location and the cell type information enables us to explore the spatial co-occurrence patterning of different cell types. We first calculated the percentage of each cell surrounded by the same type of cell and calculated the ratio of homogeneity (Fig. 5C; Methods). The results show that cancer epithelial cells, myeloid cells and B cells \ Plasmablasts were mostly surrounded by the same type of cells, while PVL, endothelial cells tended to mix with other cell types. We also calculated the spatial distance between different cell types in contact with each other based on the spatial location of the cells obtained from St2cell. The larger the value, the further the distance between the two types of cells in contact with each other, implying that their interaction relationship may be weaker (Fig. 4D). In this sample, the cancer epithelial cell-immune cell distance was overall large, suggesting a higher TME-related risk score^34^. The information established can help us to map out the immune micro-environment of cancer. Finally, to further demonstrate the results of St2cell’s reconstruction, we investigated the relationship between tertiary lymphoid structures (TLS) and annotated immune cells. Here, we selected 12-Chemokine signature in breast cancer to calculate TLS-related functions score (TLS score) of each spot, while counting the number of immune cell types cells annotated under each spot. The spatial gene expression levels of these 12 signature genes of each spot can be seen in Fig. S7. As shown in Fig. 5E, the number of annotated immune cells increased as the TLS score increased, which is compatible with the known pattern of cell type distribution in TME.

In summary, St2cell improves the spatial resolution of SRT by integrating information from histological details of co-registered images and brings new knowledge of tissue or organ architectures, enabling downstream applications such as determining the spatial distribution of cell types, depicting the spatial structure of cell populations, and mining information on spatial interactions between cells.

## Conclusion

The St2cell method integrates morphological information from co-registered HR histological images with SRT to reconstruct in situ single-cell transcriptomics. Our method combines the powers of both the deep learning model and the CQP-based model. We deploy a novel object detection model to help localize cells in tissue or organs and then utilize a contrastive learning-based model and a CQP-based model to help disentangle cell type from mixtures of mRNA transcripts at each captured location. With the fuse of HR histological images, St2cell provides a more refined depiction of the spatial patterns of cellular level gene expression profiles and identifies the spatial distribution of different cell types as well as their spatial structure in tissues. On multiple benchmark datasets, St2cell has shown superior performance in both reconstructing SRT at enhanced resolution and facilitating downstream tasks.

The steps to reconstruct in situ single-cell transcriptomics in St2cell are flexible. First, cell localization, cell feature extraction and disentanglement of the transcriptomic expressions of spots in St2cell are independent of each other, with the advantage that the reconstruction results can be further improved along with the development of computer vision and SRT technologies. Second, the CQP-based decoupling model is able to reconstruct the full genome. Unlike other image-based methods for predicting gene expression (such as Xfuse), St2cell does not require time-consuming model training for each gene, which makes it a time-efficient approach. Finally, St2cell can freely adjust the number of meta-cell types and the propensity of mixing ratios during the fitting process according to different sample situations. The flexibility of the decoupled model allows St2cell to be used for different types of SRT data.

One limitation of St2cell is that the process of cell localization and feature extraction relies on high-quality images with sufficient resolution, which to some extent limits the scope of application of St2cell, as some SRT data are currently not uploaded with raw HR images. (a common problem with current public SRT data). Another limitation of St2cell is that it assumes cells with the same histological morphology have similar gene expression, thus ignoring the intrinsic cellular variability, making the single-cell transcriptomics reconstructed by St2cell for each cell type tend to be the average expression of that cell type, resulting in some discrepancy between the gene expression distribution of real scRNA-seq and in situ single-cell transcriptomic data. To address these limitations, there is a need for retaining the highest resolved histological images during the production and preservation of SRT data, avoiding problems such as image blurring and reduced resolution due to the data compression process. Further, feature extraction methods with higher performance are needed to reduce the dependence on HR images and to better distinguish similarities and differences between cells. Integrating scRNA- or snRNA-seq over St2cell results can also alleviate the differences between the expression distribution of the two different technologies. We expect this work to be further improved to accommodate more spatial transcriptomic data and to further enhance the results.

## Methods

### Data preprocessing

To perform St2cell, we first need to preprocess the SRT data. The input data for St2cell requires high-resolution original co-registered histological images in addition to the spatial gene expression profiles. We note that the 10x Visium technology’s pipeline outputs provide a subdirectory named “spatial” which stores imaging related files. However, these images lose details of the cellular morphology as well as the background in which cells are located due to scaling. For good results, original HR histological images are needed. For spatial gene expression profiles, we performed UMI count normalization for each spot, set the total UMI count of the spot to 1e4, and then transformed to a natural log scale. Most of the analyses and validations were performed on 2,000 HVGs. However, St2cell is not limited to the number of selected genes. For application to human breast cancer, we reconstructed the full genome with St2cell. In this manuscript, we collated more than 40 samples of different tissues from 4 different data sources, including Visium FFPE adult mouse data, Visium mouse brain adjacent sections with matched snRNA-seq data, Visium FFPE human breast cancer data and ST HER2-positive tumors data as validation.

### Cell localization

In order to enhance the spatial resolution of SRT data up to cellular level, we need to accurately locate the spatial position where all cells in the sample are positioned. Here, we employ a heatmap regression-based method to achieve the cell localization in an end-to-end manner like conventional landmark localization tasks^35^. The regression step is modeled by a fully convolutional network (i.e., LinkNet^36^ in our study) with the original image as input and its corresponding heatmap as output. Each nucleus has a Gaussian-like response centered on the nucleus in the heatmap. The mean square error loss is adopted to train the model with a backpropagation strategy. Several public H&E-stained nuclei datasets (i.e., PanNuke, CoNSeP, MoNuSeg, and NuCLS) have been rectified to generate the heatmaps to train the localization model. In the inference stage, the well-trained model could output a heatmap, and the localization of all cells could be obtained by identify the local maximum responses.

### Cell feature extraction

We use a deep contrastive learning model to learn information from the histological images from spatial transcriptomic data and extract image features of cells for subsequent cell classification and cellular resolved enhancement. Here, we use the MoCo v2^37^ framework for training histological image sections. MoCo v2 is a relatively flexible framework that can obtain promising unsupervised pre-training representations on a typical 8-GPU machine with a small batch-size. St2cell uses MoCo v2 to extract image features and sets ResNet50^38^ as the backbone with a learning rate of 0.03, a batch size of 256, and an output size of 128 dimensions. Each histological image was tiled into more than 10,000 tiles with a digital resolution of 224 pixels × 224 pixels and used as training data for the contrastive learning model. For histological images with similar features from consecutive sections or the same batch, we put together whole tiles of these data for the deep learning model training parameters. For a single section, 400 epochs is the typical default parameter. For a set of multiple sections such as ST breast cancer data, we increased the training epochs to 800 due to the increase in feature information. In the data augmentation part, unlike the default settings of MoCo v2, we added additional strategies of random rotation and random vertical flip for constructing contrastive pairs. After obtaining the trained MoCo v2 model, 224 pixels × 224 pixels tiles with cells in the center are cropped and fed into the model to obtain representations of the cell image features for subsequent analysis. Here, we use to denote the image data of the i-th cell, *f*_θ_ to denote the trained deep model, and ***h***_i_ to denote the extracted feature vector. The whole process can be described as follows.

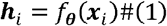

### In situ single-cell transcripomics inference

St2cell assumes that cells with similar histological morphology have similar levels of gene expression, and we classify cells based on their image features and use this as the basis for assigning meta-cell types to each cell. For spots, here we assume that the expression level of a spot can be obtained by averaging the expression levels of the cells covered under the spot. Different spots have different composition ratio of meta-cell types and different gene expression levels. Therefore, it is possible to relate the gene expression of meta-cell types to that of spots, thus solving the reconstruction of gene expression profiles of each cell and achieving super-resolved enhancement of the spatial transcriptome. This is a typical weakly supervised problem where the optimization goal is how to assign values to the gene expression of each meta-cell type so that their combined expression level is closest to the known measurement of the spot.

We first classified the cellular image features obtained from the deep contrastive model, i.e., the feature vectors are classified into m classes using KMEANs, as follows.

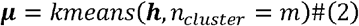

Further, for the i-th cell and the j-th cluster center, we calculated the cosine distance between them. Here we refer to the strategy of t-sne^39^ and use the Student’s t-distribution as a kernel to measure the similarity between cells and cluster centers as a soft assignment.

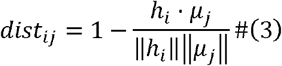

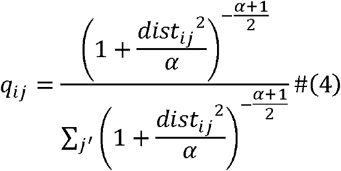

After obtaining the distribution ***q***_*i*_ of the i-th cell with respect to all cluster centers, we normalized the distribution of each cell to obtain the probability ***p***_*i*_ that each cell is composed of different meta-cell types or clusters. We compute the target distribution ***P*** using softmax as follows, where *τ* is a temperature hyperparameter to adjust the target distribution. Here, to make the cluster centers easier to interpret, we prefer to choose a larger *τ* such that each sample consists primarily of one meta-cell type as much as possible.

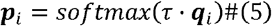

From the above calculations, we obtained the meta-cell type composition for each cell. The coordinate information of the captured location and cells was further combined to calculate the probability of the meta-cell type constituting each spot. Here we set the cells contained in the same spot to have the same weight.

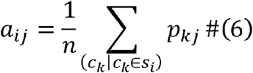

In which, *c_k_* is the k-th cell and *S_i_* is the i-th spot. We use pixel distance to denote the Euclidean distance between the center of the spot and the cell, as follows.

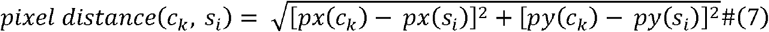

Where (px, py) denote the coordinates (in pixels) of the cell or spot in the image. For the i-th spot and the k-th cell, *C_k_* ∈ *S_i_* if the distance between them is less than a given threshold. Considering the spread of staining between spots, St2cell sets the default threshold to the size of the spot’s diameter. The probability distribution ***A*** of spot consisting of different cell categories was obtained by Equation 6 (n equals the size of *S_i_*).

The SRT data from ISC technologies contain spot-level gene expression profiling, while the gene expression level of different cells is unknown. For a particular gene, we use *y_i_* to denote the known expression level of this gene in the i-th spot and the parameter *w_j_* to learn the expression level of this gene in the j-th meta-cell type. To solve the problem of non-negative expression levels ***w*** of genes of different meta-cell types, we minimize the mean square error (MSE) between the known gene expression levels of spots and the average of the expression levels of all meta-cells contained in the spots. It’s a constrained optimization problem for which a MSE loss is to be minimized subject to constraints ***w*** ≽ **0**. Also, considering the distinctive feature of scRNA-seq data is the increased sparsity, i.e., fraction of observed “zeros”, we added the *l*1 sparse regularization ***w*** of as a extra constraint and used λ to adjust the ratio of the sparse regularization to the MSE (0.2 was used as our default parameter). Thus, the optimization problem can be written as:

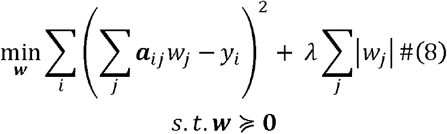

The existence of sparse constraints makes the form of the above optimization problem difficult to solve. Therefore, we solve the equivalent problem of the original problem, i.e., we add the variable ***v*** same number of elements as ***w*** and add new constraints, so that the original optimization problem is modified as:

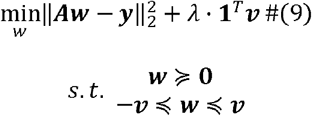

Then, we reformulate the problem into a quadratic programming problem:

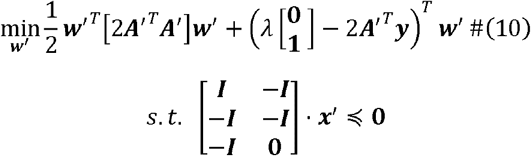

Where ***A′*** = [***A,0***], ***w′*** = [***w, v***]^*T*^. Since ***A′^T^ A′*** is semi-positive definite, this problem is a convex quadratic programming problem with a globally optimal solution. Here, we use the CVXOPT, a free software package for convex optimization based on the Python programming language, to solve this problem. After calculating the optimal solution ***w***, we calculate the reconstructed gene expression levels of the i-th spot and cell, respectively, as:

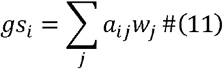

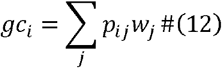

### Experimental settings

#### Visium brain data processing

The Visium FFPE adult mouse brain data contains 3,000 spots. We set the number of meta-cell type to 500, and the diameter threshold to 188 pixels, and clustered over 30,000 cell image features. After obtaining the reconstructed expressions at the cellular level, we clustered these expressions of cells into 20 categories. Based on the co-registered histological image and known spatial gene expression profiling at captured locations, we manually labeled the boundaries of the anatomical regions.

Visium and snRNA-seq from adjacent sections in the mouse brain from Kleshchevnikov et al.^18^ were also used. This data has 5 adjacent sections with approximately 3000 spots each. The settings of St2cell are consistent with previous parameters for processing adult mouse brains, except for a diameter threshold of 144 pixels. Unlike the previous adult mouse brain data, this data was accompanied by snRNA-seq. After obtaining cellular resolved expression profiles for each section with St2cell, we integrated the extra single-nucleus data.

#### ST Breast cancer processing and ARI calculation

Since the number of spot per sample for ST breast cancer data is around 500, we used only 250 pseudo-cell types to avoid getting ill-conditioned algebraic systems. The same default parameters were used for all samples. For the cell-level expressions obtained from St2cell reconstruction, we used PCA to downscale to 50 dimensions, and then clustered according to the number of manually labeled categories. For “G2” and “H1”, we compared the clustering result of St2cell with Kmeans, Louvain, SpaGCN and BayesSpace when the number of clusters was set at seven for all methods. ARI is used to evaluate the results of spatial domain discovery.

#### Visum breast cancer processing

For Visium FFPE breast cancer data, we set the number of meta-cell type to 500, and the diameter threshold to 188 pixels. Unlike previous treatments, we here reconstructed all gene expression at the cellular level as query data. Next, we screened the genes that co-occurred in the reference and query data, and removed other genetic data that did not overlap. We used the filtered data as input data for Scanpy API “scanpy.tl.ingest” ^40^ and obtained cell annotations.

#### Settings of spatial domain detection

Spatial domain detection is actually a clustering problem. Traditional clustering methods, such as Kmeans and Louvain, take only gene expression data as input, while clustering methods specifically designed for SRT, such as SpaGCN and BayesSpace, take spatial information into account. While St2cell goes a step further and uses deep learning to further integrate spatial as well as image information. We first preprocess the gene expression at each spot in the SRT using different methods and then perform clustering after obtaining the characteristic expression at the captured location. Where the number of categories is aligned with the number of manually annotated categories included in that sample. Finally, we use ARI to calculate the agreement of the spatial domain detection results with the real annotations. Kmeans and Louvain are implemented with “stLearn”^41^, and the officially recommended parameters are set as running parameters. SpaGCN and BayesSpace are also run with default parameters. Unlike the other methods, St2cell first obtained in situ single-cell transcriptomics data and selected the reconstructed expression levels of 2,000 HVGs as cellular features, and if St2cell had the same parameters as the other methods, the two methods were aligned on this parameter. Finally, the cell expression levels contained in each spot were averaged according to the spatial location of the cells and spots, and the ARI was calculated after clustering the obtained spot expression features.

#### Blurred ST data generation

To test the performance of st2cell for resolution enhancement and resistance to noise, we first designed an input data processing strategy to generate blurred data. Here, we simulated the contamination of spot’s gene expressions by surrounding spots using ST breast cancer data. For each spot of the sample, we averaged the gene expression levels of its surrounding spots using a 3×3 kernel to obtain the blurry expression profiles of the corresponding captured locations. By the above operation, we can get a new SRT sample with noise expression profiling. For St2cell, the cells contained in each spot of the new blurred sample are the sum of the cells contained in the corresponding 9 spots of the original sample. We used St2cell to obtain cellular resolved gene expression profiles and recovered gene expressions of spots from noise-contaminated capture locations based on the actual cells contained under each spot. For BayesSpace, we used it to enhance the resolution of the blurred samples. For each spot of the original sample, we averaged the gene expression levels of the 9 corresponding sub-spots on the BayesSpace-enhanced sample as the recovered result.

#### Lower-resolution ST data generation

To test the accuracy of St2cell reconstructed expression levels, we did a more rigorous input data processing to reduce the spatial resolution of the ST data, and also used BayesSpace as the baseline for comparison. Here, we synthesize 9 spots from every 3 rows and 3 columns of the original data into one spot and give it to the generated new data. The coordinates of the newly synthesized spot are the center coordinates of the original 9 spots. As the way we operated before, we added the expression levels of the 9 spots as the expression values of the new spot. Moreover, in order to obtain a lower resolution, each spot in the original data is involved in the synthesis process only once. After the ST breast cancer data was reduced to 9 times the resolution of the original data, the number of spots was roughly around 50. Thus, we set the number of meta-cell types in St2cell to be reduced to 25 accordingly. For St2cell, the cells contained by each new spot are all the cells contained by the corresponding 9 spots in the original sample. Therefore, we can still obtain the reconstructed expression profile at cellular resolution and are able to calculate the gene expression level of the deleted spot to recover the original resolution. For BayesSpace, it can enhance the samples to 9 times the resolution of the input data, and the enhanced samples have exactly the same resolution as the original samples.

For the generated blurry and low-resolution ST data, we calculated the correlation coefficients between the reconstructed expression levels of the spots obtained by both methods and the known expression levels of the original samples as a performance indicator to evaluate the enhanced resolution of the methods.

#### SnRNA-seq integration

As with PRECISE^33^ and SpaGE^30^, we first projected the reconstructed cellular resolved expression profiles and real single-nucleus expression profiles into the common latent space separately, and applied independent Principal Component Analysis (PCA) on the two expression matrices to obtain their top 50 principal components separately. Then, the cosine similarity matrices of these independent components are calculated and SVGs decomposition is performed. The decomposition results are used to align their principal components, and the expression matrices projected into the same latent space are calculated based on the centers of the principal vectors obtained. Subsequently, the integrated results are obtained by weighting the single-nucleus data with the KNN approach based on the cosine distance of the principal vectors. For more details of the calculation, please refer to the relevant section of SpaGE.

#### Annotating in situ single-cell transcriptomic maps

A Visium FFPE human breast tissue from BioIVT Asterand Human Tissue Specimens annotated as “ductal carcinoma in situ, invasive carcinoma” was used here (see also Data availability). For annotating cellular resolved transcriptomic maps obtained by St2cell, we also downloaded scRNA-seq data of breast cancer generated by Wu et al. through the Broad Institute Single Cell portal as an appropriate set of prior biological knowledge^42^. The annotated results (or the “query” dataset) were then compared to the expert-annotated reference dataset we downloaded earlier, and the labels were transferred from the reference cell to the sufficiently similar cell in the query dataset. The Scanpy API “scanpy.tl.ingest” was used for automatic cell annotation.

#### Calculation of metrics used in tumor microenvironment experiment

For inferred contacts of cell types within Visium human breast cancer sections, we used the Delaunay triangle method to construct graphical structures whose edges represent contacts of the annotated cell type. TLS are ectopic lymphatic formations with structure that containing B-cell follicles and germinal centers surrounded by T-cell regions, which form within non-lymphoid tissues^43^. As for the calculation of TLS score, we selected CCL2, CCL3, CCL4, CCL5, CCL8, CCL18, CCL19, CCL21, CXCL9, CXCL10, CSCL11, CXCL13 in breast cancer as confirmed by previous studies^44^, and the expression levels of these genes were summed to calculate the TLS score.

## Data availability

The following datasets were used in the evaluation of St2cell. These datasets can be obtained from the following websites or accession numbers: (1) Visium FFPE Adult Mouse Brain (https://www.10xgenomics.com/resources/datasets/adult-mouse-brain-ffpe-1-standard-1-3-0). (2) Visium Mouse Brain and snRNA-seq from adjacent sections in the mouse brain (ArrayExpress E-MTAB-11114, E-MTAB-11115 and https://github.com/BayraktarLab/cell2location/). (3) Visium FFPE Human Breast Cancer, section from invasive ductal carcinoma of human breast (https://www.10xgenomics.com/resources/datasets/human-breast-cancer-ductal-carcinoma-in-situ-invasive-carcinoma-ffpe-1-standard-1-3-0). (4) ST Breast Cancer, sections from HER2 positive breast tumors (https://github.com/almaan/her2st)

## Code availability

An open-source implementation of the St2cell algorithm can be downloaded from https://github.com/Svvord/St2cell.

