## Supplementary for "St2cell: Reconstruction of in situ single-cell spatial transcriptomics by integrating high-resolution histological image"

**Supplementary information**

**Supplementary Fig.S1 Visualizations of the expression distribution of six regional-enrichment markers – Satb2 (CTX), Rgs9 (L4), Prox1 (L5a), Aldh1a1 (L5), Mbp (L6), and Gpr165 (L6b) obtained from in situ hybridization data.**

**Supplementary Fig.S2 Visualizations of the expression distribution of eleven layer-specific markers obtained from in situ hybridization data.**

**Supplementary Fig.S3** **Visualization of the expression distribution of the L6b marker, Cplx3, the high expression position of this marker can be used as a demarcation line between WM and CTX. (A)** St2cell’s reconstruction of Cplx3 expression levels consistent well with histological image features, with arrows pointing to the WM region sandwiched between CTX and HPC. **(B)**. ISH data of Cplx3, with arrows pointing to WM. **(C)** ISH technique obtained for the expression distribution of Cplx3.

**Supplementary Fig.S4 Reconstructed Epcam expression level of “H1” by St2cell and BayesSpace under two different simulations.**

**Supplementary Fig.S5 St2cell has a certain level of robustness in terms of utilizing noisy images feature.**

**Supplementary Fig.S6 Manual annotation of Visium FFPE Human Breast Cancer.**

**Supplementary Fig.S7 Spatial gene expression levels of 12-Chemokine signature of each spot.**

**Supplementary Fig. S1**

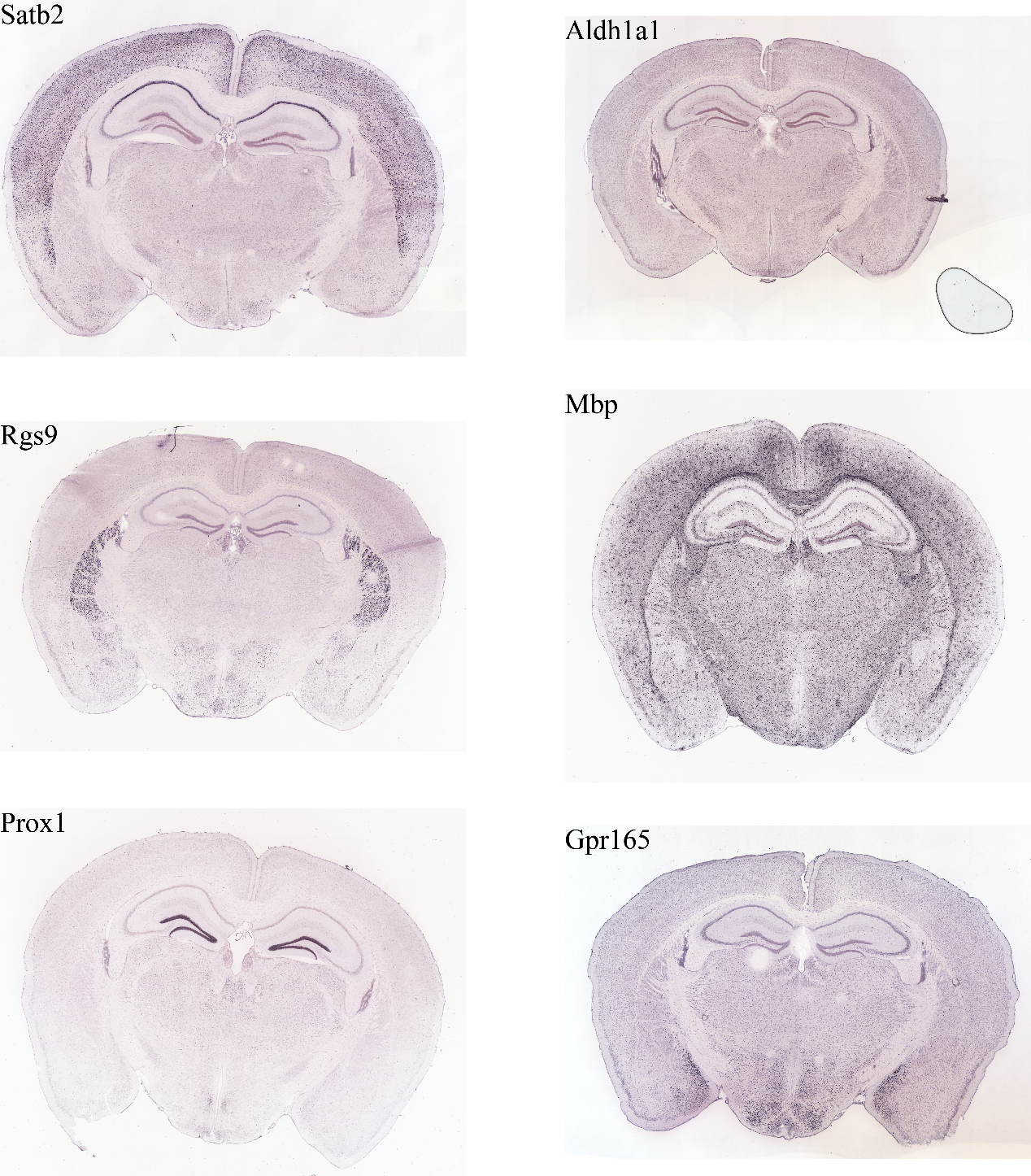

**Supplementary Fig. S2**

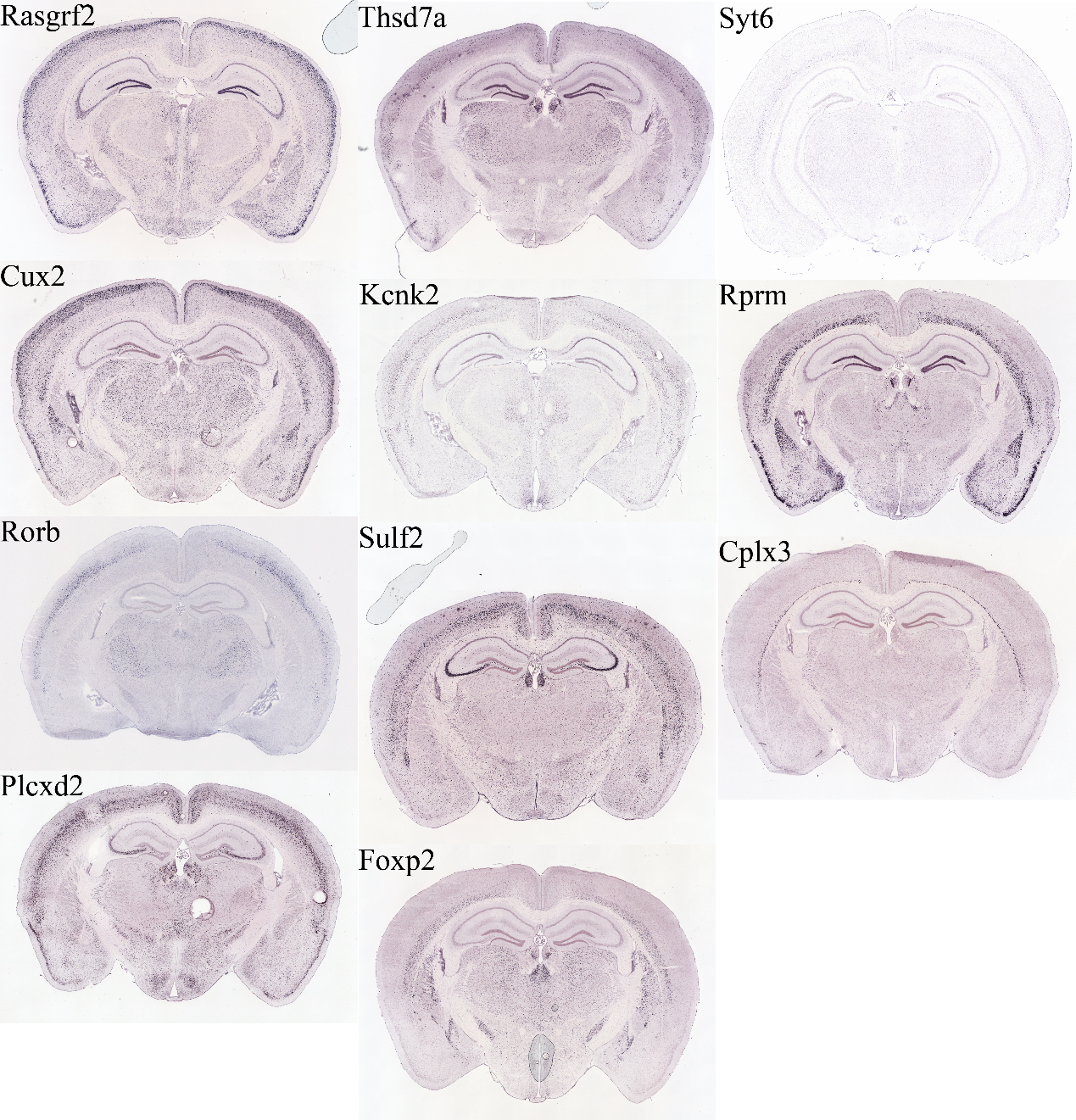

**Supplementary Fig. S3**

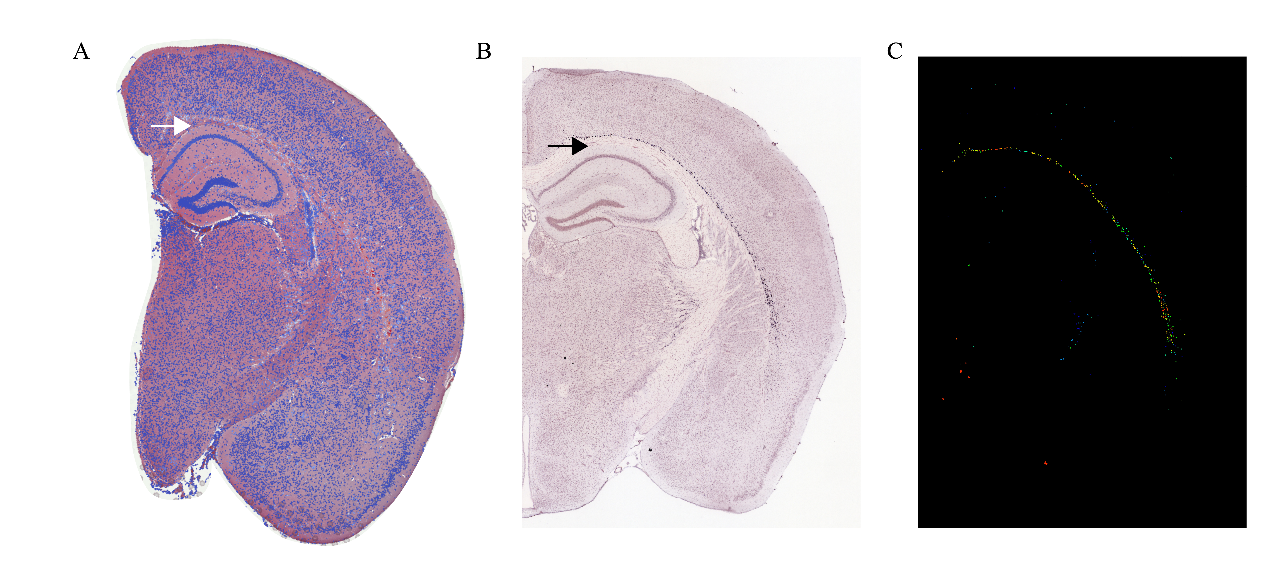

**Supplementary Fig. S4**

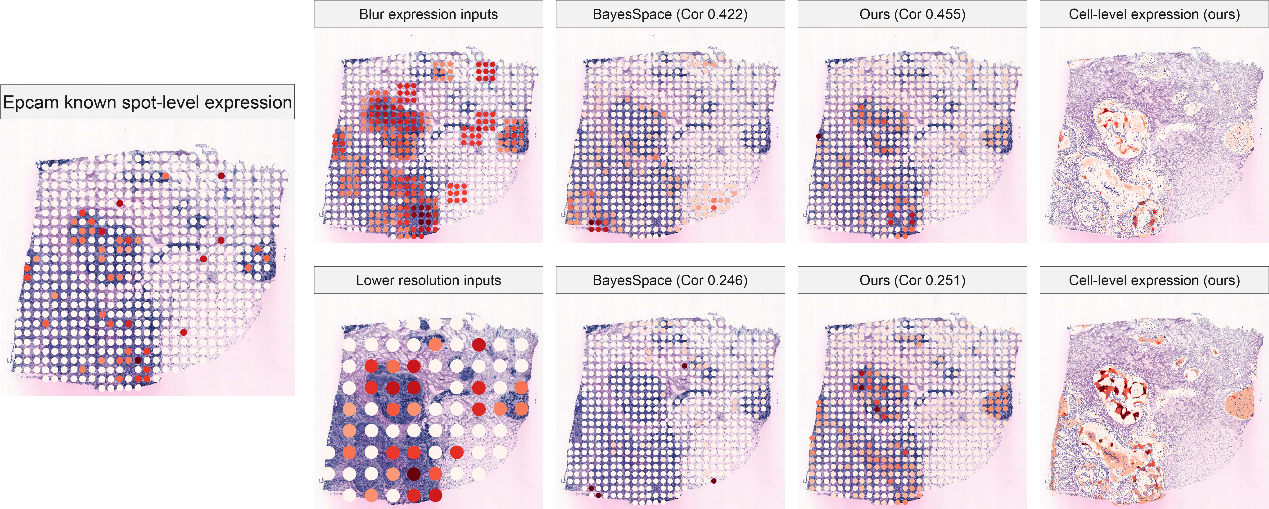

**Supplementary Fig. S5**

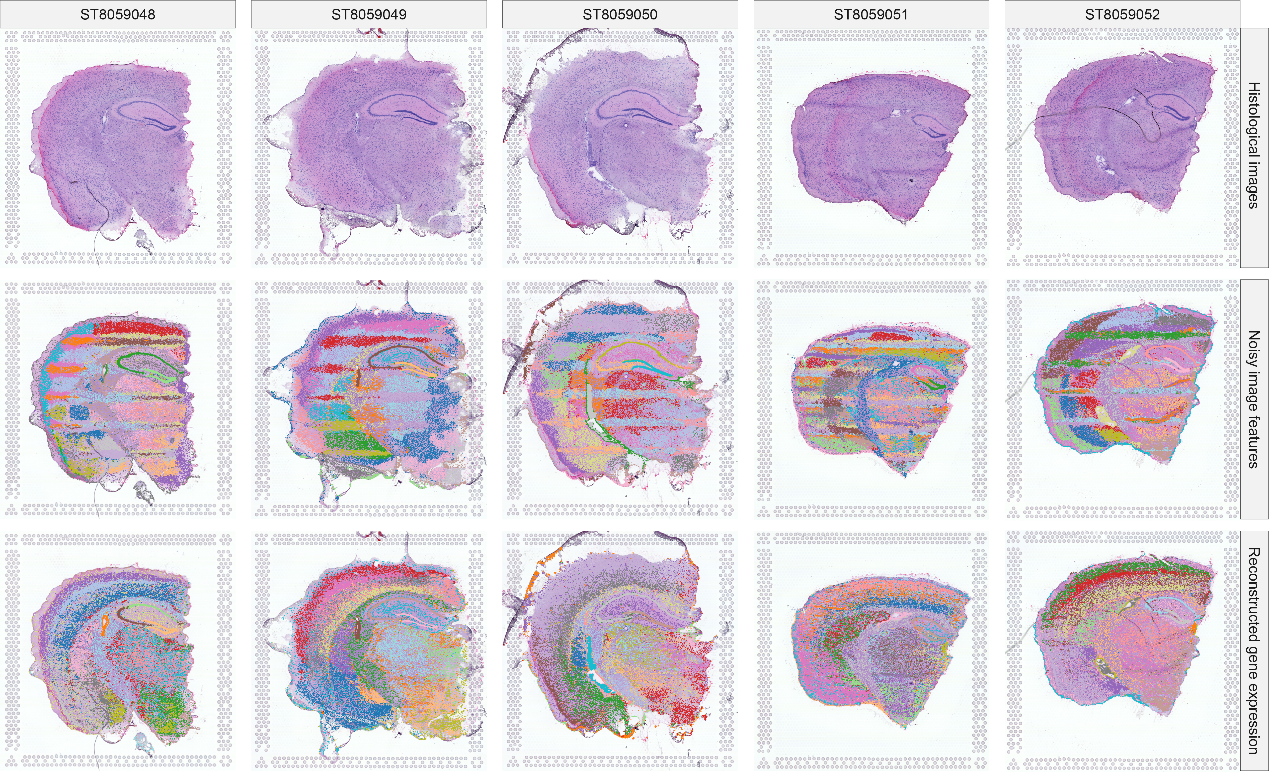

**Supplementary Fig. S6**

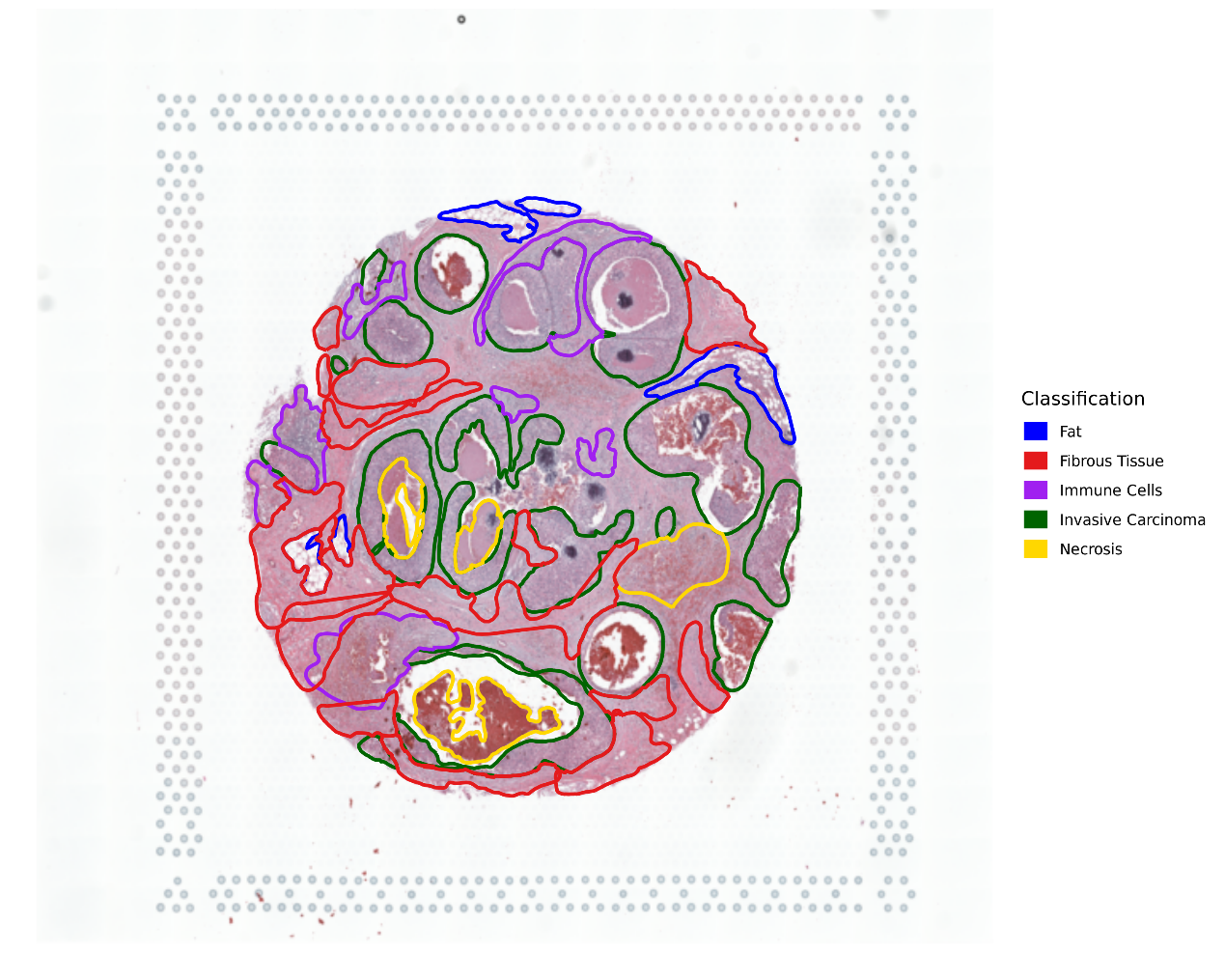

**Supplementary Fig. S7**

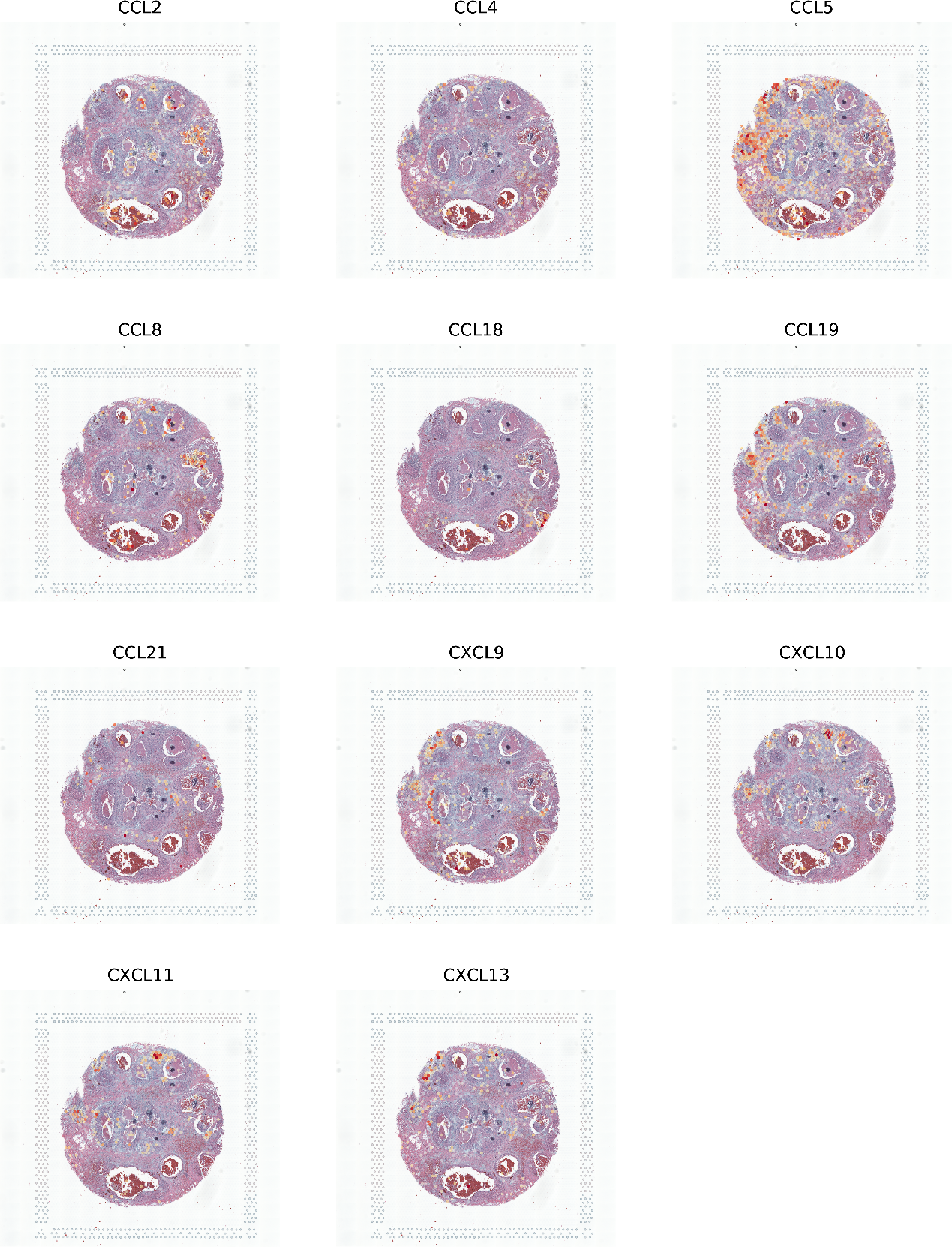
